# Construction of a drug sensitised *Candida auris* strain

**DOI:** 10.1101/2024.12.28.630619

**Authors:** Mahnoor Hassan, Jessica Furner-Pardoe, Yiyuan Chen, A. James Mason, Khondaker Miraz Rahman, J. Mark Sutton, Charlotte Hind, Barry Panaretou

**Affiliations:** Institute of Pharmaceutical Science, King’s College London, 150 Stamford Street, London. SE1 9NH, United Kingdom; Vaccine Development and Evaluation Centre, UK Health Security Agency, Porton Down, Salisbury, Wiltshire, SP4 0JG, UK; Klura Labs, The Epicentre, Haverhill Research Park, Enterprise Wy, Haverhill CB9 7LR, UK

**Author notes:** To whom correspondence should be addressed: Barry Panaretou, Institute of Pharmaceutical Science, King’s College London, 150 Stamford Street, London, UK, SE1 9NH., Charlotte Hind, Antimicrobial Discovery, Development and Diagnostics, Vaccine Development and Evaluation Centre, UK Health Security Agency, Porton Down, Salisbury, Wiltshire, UK, SP4 0JG.

**Keywords:** *Candida auris*, drug efflux pumps, drug screening

## Abstract

*Candida auris* is an emerging pathogenic fungus causing outbreaks of invasive disease in healthcare facilities worldwide with the majority of clinical isolates demonstrating intrinsic resistance to multiple drug classes. Therefore, there is a pressing need for new antifungals, but screening in wild-type *C. auris* strains may miss scaffolds which would be good subjects for med-chem projects as antifungals. Work presented here indicates that deletion of *CDR1* and *MDR1* in *C. auris*, respectively encoding an ATP-binding Cassette and a Major Facilitator Superfamily efflux pump, generates a strain that is hypersensitised to drugs and can act as a platform for antifungal drug screening. The *cdr1*Δ*mdr1*Δ mutant was 64- and 8- times more sensitive to fluconazole and voriconazole, respectively, compared with the wild-type strain. In a pilot study using a 2,000 compound library, use of the *cdr1*Δ*mdr1*Δ mutant identified over twice as many compounds compared to wild-type. When assessing the capacity to form biofilms, there was a modest difference in biomass between the wild-type and *cdr1*Δ*mdr1*Δ mutant, but there was little to no statistically significant difference observed when using a colorimetric dye that measured metabolic activity. Little to no statistically significant difference was observed between wild-type and the efflux pump mutant in virulence, as judged by the larval *Galleria mellonella* infectivity assay. Accordingly, the *cdr1Δmdr1Δ* mutant will be useful when screening for drugs that limit formation of biofilms or limit fungal virulence, as these two properties are unaffected when drug efflux is compromised.

## 1. Introduction

Initially reported in Japan in 2009, *Candida auris* is an emerging multi-drug resistant (MDR) fungus associated with an increase in nosocomial outbreaks across five continents with reports of occurrences across more than 20 countries [1–3]. Six distinct geographical clades are recognized: Clade I (South Asian, India and Pakistan), clade II (East Asian, Japan), clade III (African, South Africa), clade IV (South American, Venezuela), clade V (Iran) and clade VI, reported in 2023 from strains isolated in Singapore [2, 4–6].

Whole-genome sequencing (WGS) of isolates revealed that the clades diverge from each other by tens of thousands of single-nucleotide polymorphisms (SNPs) [3, 4]. Within each clade, isolates appear highly related, differing by less than a thousand SNPs [3, 7]. This appears to suggest *C. auris* emerged as a human pathogen almost simultaneously, and perhaps independently, in the geographical clades [3]. Within different clades, variations of virulence have been observed, with isolates from clade IV presenting the highest mortality rate in silkworm infection models (∼80% after 21 hours) [8]. Furthermore, resistance to current fungal treatments also appears to have observable variations, with clade II reportedly presenting less drug resistance than the other clades [9, 10].

Characteristics that separate *C. auris* from its closest relatives include the ability to grow at 42°C, carbon source assimilation, limited ability to form hyphae and, importantly, its resistance to multiple antifungal drugs [11]. *C. auris* has an incredibly high mortality rate of up to 60% in immunocompromised patients [3, 12]. Clinical occurrence of *C. auris* has rapidly progressed from minor cases of superficial infections to extremely invasive bloodstream infections [13]. A study carried out by Navarro Arias *et al*., 2019, indicated that *C. auris* was able to evade key aspects of the human immune response, such as phagocytosis by macrophages and neutrophil-mediated cell death, failing to induce an adequate inflammatory response [14]. *C. auris* also has an extraordinary capacity to colonise and proliferate on surfaces, exhibiting the ability to withstand higher temperatures than other *Candida* spp. and exposure to oxidative stress by hydrogen peroxide, cell wall stress and cationic stress, resulting in a greater resistance to disinfection procedures [15, 16]. Therefore, the lack of optimal therapeutic interventions against *C. auris,* due to its reduced susceptibility to antifungals present in clinical settings, further contributes to the poor, often fatal, outcomes amongst infected individuals. In fact, *C. auris’* multi-drug resistance capacity is arguably the most concerning aspect of its pathology. Many clinical isolates of *C. auris* are resistant to the most commonly used drugs, the azoles, whilst so*me C. auris* isolates are pan-drug resistant to all classes of antifungal drug, including the echinocandins, the latest class of antifungals to reach the clinic [17, 18].

Discovery of new drug candidates against *C. auris* is hindered by the over-expression of ABC transporters in this species, which can efflux out any compound which is a substrate for these efflux pumps [19] . Two genes that are significantly over-expressed in several drug resistant isolates of *C. auris* are the efflux pumps *CDR1* and *MDR1* [19, 20]. Whilst single efflux pump knockout strains of *C. auris* (*cdr1Δ* and *mdr1Δ*) have been created before [20] we have created the first example of a double efflux pump knockout (*cdr1Δmdr1Δ*) in *C. auris*. We created both single (*cdr1Δ* and *mdr1Δ*) and double (*cdr1Δmdr1Δ*) knock-out *C. auris* strains, without significant impact on virulence or biofilm forming capability. The *cdr1Δmdr1Δ* strain was used to screen a 2000 compound library, where higher numbers of drug candidates were identified by the double knock-out strain compared to the parent strain.

## 2. Materials and Methods

### 2.1 Strains and Growth conditions

*Candida auris* strain TDG1912 is a clade III strain, isolated from air in a hospital ward. Yeast strains were grown in either Yeast extract Peptone Dextrose (YPD) or standard defined (SD)/minimal media [21] or RPMI-1640 MOPS buffer + 2% glucose (RPMI 2% G) at 37°C, with constant shaking at 200 rpm. For solid YPD/ SD media, 2% agar was added.

### 2.2 Gene modification cassette construction

The *NAT1-Clox* deletion cassette, described by Shahana *et al*., 2014 [22] was used to allow reuse of the *NAT1* selective marker gene *via* its excision using recombination across *lox*P sites mediated by Cre recombinase. Only one round of transformation was necessary *per* gene deletion, because *C.auris* is haploid [23]. For the construction of a gene-specific deletion cassette, PCR was used to amplify the *NAT1*-*Clox* sequence, from pLNMCL [22] and ∼500 bp DNA sequences upstream and downstream of the *C.auris* gene of interest. The separate fragments were combined within pSL1183, a derivative of pSL1180 [24] using HiFi DNA Assembly (New England Biolabs, USA). Deletion cassettes composed of the 5’ flanking homology domain, *NAT1-Clox*, and the 3’ flanking homology domain were amplified using Q5® High-Fidelity DNA polymerase (New England Biolabs, USA). Sequences of oligonucleotides required for generating all amplicons, are provided in the Supplementary file S1.

### 2.3 Transformation of gene modification cassette into *Candida* spp

To prepare cells for transformation *C,auris* was inoculated into YPD and grown to exponential phase (5 × 10^6^ to 8 × 10^7^ cells/mL) overnight at 30^0^C with agitation (140rpm). A modified version of the method described by Kim *et al*. was used to render cells electrocompetent [25]. Briefly, cells were pelleted and resuspended in 40 mL of LiACTE Buffer (0.1 M Lithium acetate, 10 mM Tris-HCl pH7.5, 1 mM EDTA) and incubated at 30°C. After 1 hour, 1 mL of 1 M dithiothreitol (DTT) was added to the cells and further incubated for 30 minutes at 30°C. Cells were washed once in ice-cold sterile dH_2_O followed by a wash in ice-cold 1 M Sorbitol. Cells were resuspended in a volume of ice-cold 1 M Sorbitol sufficient to yield a cell density of 1 × 10^8^ cells/mL. The gene-specific cassette (3 μg, prepared as described previously [25]) was added to 40 μL electrocompetent cells and electroporated in a Bio-Rad Pulse Controller-Gene Pulser™ (1.8 kV, 200 Ω, 25 μF) followed by addition of ice-cold 1 M Sorbitol to give a final volume of 1 mL. Cells were pelleted, resuspended in 10 mL of YPD and incubated at 30°C with agitation (140 rpm). After 4 hours cells were pelleted and resuspended in 400 μL of fresh YPD, and inoculated across four YPD agar plates containing nourseothricin (200 μg/mL), supplemented with 2.5 mM cysteine and 2.5 mM methionine (to fully repress Cre recombinase-mediated excision of the *NAT1* drug resistance marker), and incubated at 30°C for 2 days. When sub-cultured, transformants were maintained on the same media. Genomic DNA was extracted using the YeaStar^TM^ Genomic DNA extraction kit (Zymo Research, USA) and used as template for diagnostic PCR to confirm locus-specific integration of the deletion cassette.

The deletion cassette includes the *Cre* recombinase gene under the control of the *C.albicans MET3* promoter. Incubation on SD media (i.e. in the absence of methionine and cysteine) is supposed to lead to excision of the deletion cassette *via* Cre recombinase-mediated recombination across *loxp* sites, but this was inefficient in our hands. Instead, the *NAT1*-Clox cassette was removed as described by Reuß *et al*., 2004 [26]. Cells were incubated in SD broth for 6 hours at 30°C, 140 rpm and inoculated onto YPD agar (100-200 cells per plate) supplemented with a sub-lethal concentration of nourseothricin (25 μg/mL) and grown for upto 4 days at 30°C. Nourseothricin sensitive cells, generated by excision of the *NAT1* selective marker gene, would grow comparably slower than nourseothricin resistant cells, giving rise to smaller colonies. In this way, many thousands of colonies could be screened, in order to identify those arising from a cell in which recombination had occurred across the *lox*P sites.

### 2.4 Primers for diagnostic PCR

CDOUTF2 GAGGTTCCAAGGCCGGACAACGTGCG

CDOUTR CTGGCTCGTTGTTGTAATTGATCCCACATT

CdrFor ATGTCCGAGAAACCTTTTGTCGACG

CdrRev GGTACCGATCATGTTGATAGCGATG

MdrF2 ATAATCCGTGTACGAGGCGTGTTTC

MdrR3 TCAAAGTTGTGTACAGTGAGCGGAG

WCLOXF2 GATACTTGGCTTGGTCTGGTCACTCTGC

WCLOXR TATTAGTACTGGATCTATAACTTCG

### 2.5 Galleria mellonella infection assay

All assays were performed on 10 larvae (n=3, 30 larvae in total) *per* condition. Healthy larvae were used within 2 weeks of receipt from the supplier (Livefoods Ltd, UK). *C. auris* cells were picked from a YPD agar plate into 3 mL of YPD broth and grown overnight at 37°C with shaking. The following day cells were diluted to OD_600_ 2.0 in PBS. 10 μL of cells (∼2 x 10^5^ cells) were injected through one of the prolegs of each larva. Wild-type *C.auris* TDG1912 in PBS and PBS-only were used as positive and negative controls, respectively. Survival of the larvae at 37°C was observed for 72 hours. Statistical analysis was performed using GraphPad Prism, utilising Log rank (Mantel Cox) and Gehan-Breslow-Wilcoxon tests on Kaplan-Meier survival curves.

### 2.6 Crystal violet biofilm analysis

Cells were backdiluted from an overnight culture to an OD_600_ of 0.01 in RPMI 2% G, 200 µL/well was added to two columns on a 96 well plate and the plate was incubated at 37°C for 24 hours. Planktonic cells and liquid media were then carefully removed from the wells and biofilms were washed once with 200 μL PBS. The biofilm was fixed for 60 minutes at 80°C in a heating block then stained using 200 μL 0.05% crystal violet (CV) for 15 minutes at room temperature. CV was then discarded, the plate was washed with dH_2_O and allowed to air dry under a sterile laminar flow. The biofilm was de-stained with 200 μL of absolute ethanol for 20 minutes, after which absorbance readings were obtained at OD_570_ for each well. Data was normalised using the following formula: (Sample – Average negative control) ÷ (Average positive control – Average negative control). Statistical analysis was performed using GraphPad Prism, utilising t-tests (unpaired, parametric) on normalised data from three biological replicates, with 16 technical replicates each (n=48).

### 2.7 XTT metabolism biofilm analysis

This method was modified from a protocol for assessing biofilm formation in *Candida* spp. developed by Pierce *et al*., 2008 [27]. The overnight cell culture was pelleted and washed twice in ice-cold PBS and diluted to a cell density of 1 × 10^6^ cells/mL in RPMI 2% G. Each strain was aliquoted across two columns of wells on a 96 well plate (200 μL of cells in RPMI, in each well) followed by incubation at 37°C for 24-48 hours. Planktonic cells and liquid media were carefully removed and wells were washed once with 200 μL PBS. Before removing the PBS, absorbance readings at OD_490_ were obtained for the plate, after which the PBS was carefully discarded. XTT mixture (1 μL of 10 mM menadione in 10 mL 0.05 %w/v XTT [in PBS]), was aliquoted at a volume of 100 μL into each well. The plate was incubated in the dark for 3 hours at 37°C. After the incubation, absorbance readings at OD_490_ for each well were recorded. Data was normalised and statistically analysed as described for the crystal violet method.

### 2.8 Drug susceptibility (MIC) assay for *Candida* spp

Minimum inhibitory concentrations were determined, following EUCAST guidelines [28]. A high concentration stock solution of compounds was made up in DMSO. The compounds were diluted in RPMI 2% G, ensuring the final DMSO concertation was below 2%, as this concentration of DMSO did not impact cell growth. A serial dilution (1:2) of drugs/compounds (100 μL per well) was prepared across a row in 96 well plate. RPMI 2% G only (without drug) was aliquoted into the final column as a positive control for the growth of cells. Cells were backdiluted from an overnight culture to an OD_600_ of 0.01 and 100 μL/well was added to the plate. The plate was incubated for 24 hours in a static incubator at 37°C, after resuspension of cells, absorbance readings were obtained at OD_530_. Relative cell growth in the presence of the compound was compared against a positive control for growth to deduce MIC_90_ (amphotericin-B) and MIC_50_ (azoles, echinocandins and azole-scaffold drug candidates) values, with modal averages calculated for three biological replicates. Both MIC_50_ and MIC_90_ were calculated for dequalinium chloride, as no pre-existing information on which readout is relevant is available for this compound.

### 2.9 High-throughput screening of the PrimScreen2 library

A 2000 drug-like compound library (Primscreen2) was purchased from OTAVAChemicals (Lithuania; https://otavachemicals.com/sdf). The compound library was screened against the *cdr1Δmdr1Δ* and the *C.auris* TDG1912 parent strain, at a single concentration of 10 µM in 96 well plates using a CyBio Felix liquid handler. The DMSO concentration was kept below 2%. Susceptibility was determined as above for MICs.

## 3. Results

### 3.1 Drug sensitised strain construction

Double deletion of *CDR1* and *MDR1* was performed on fluconazole resistant *C. auris* isolate TDG1912. This mutant was generated by initially deleting *CDR1* by replacing the gene with a targeted deletion cassette containing a *NAT1* selective marker. Rates of targeted integration in *C.auris* are low [29], necessitating analysis of large numbers of transformants. For this reason nourseothricin resistant transformants were transferred to YPD solid media supplemented with 50 μg/mL cyclohexi(i)mide because drug-pump deletes are hypersensitive to this drug [30]. Seven of the nourseothricin resistant transformants are shown in Figure 1A, with two of them being hypersensitive to cyclohexi(i)mide (Figure 1A, sectors 2 and 6). Diagnostic PCR indicated that of these two, only one of them (corresponding to the transformant tested in Figure 1A, sector 6) was a *CDR1* delete (Figure1B, C and D). Amplification across *CDR1* could not be used as a diagnostic reaction because the size of this locus is similar to that of the deletion cassette that was designed to replace it. For this reason, integration at the targeted site was identified by amplification of the 5’ and 3’ junctions formed by homologous recombination of the deletion cassette at *CDR1*. Both junctions were amplified from one of the transformants that was hypersensitive to cylohexi(i)mide, a 969bp amplicon at the 5’ junction (Figure 1B, lane 6) and a 1389bp amplicon from the 3’ junction (Figure 1C, lane 6). As expected, a 4470bp region within *CDR1* could not be amplified from this transformant (Figure 1D, lane 6) and was amplified from the wild-type (Figure 1D, lane W+).

**Figure 1.**
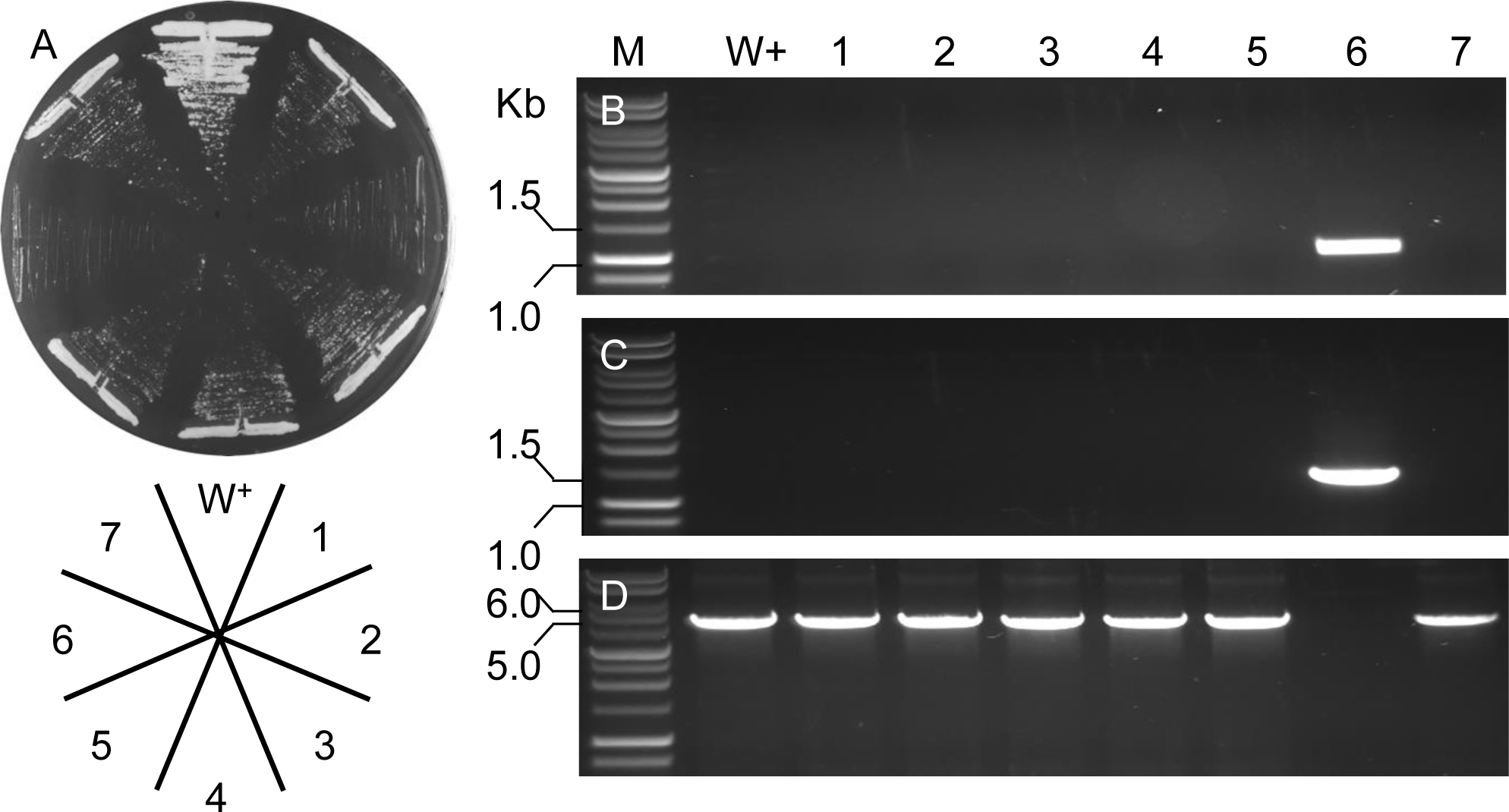
*CDR1* gene deletion in *C. auris*. Wild-type *C.auris* was transformed with a cassette designed to replace *CDR1.* (A) Nourseothricin resistant transformants were transferred to YPD + cyclohexi(i)mide (50μg/ml). Seven of the transformants that were sensitive to cyclohexi(i)mide are shown here. Diagnostic PCR using genomic DNA extracted from wild-type (W+) and transformants tested in (A) using oligonucleotide primers (B) CDOUTF2/WCLOXR, (C) WCLOXF2/CDOUTR and (D) CdrFor/CdrRev (C). Sizes of markers (M) are indicated on the left.

*MDR1* was also deleted in wild-type *C. auris*. Once again nourseothricin resistant transformants were tested for sensitivity to cyclohexi(i)mide compared to wild-type, of the eight transformants tested, one was sensitive to cyclohexi(i)mide (Figure 2A, sector 6). Diagnostic PCR across *MDR1* indicated targeted deletion of this gene as an amplicon corresponding to the deletion cassette (5965bp, Figure 2B, lane 6) was generated as opposed to an amplicon corresponding to intact *MDR1* (2966bp, Figure 2B, lane W+).

**Figure 2.**
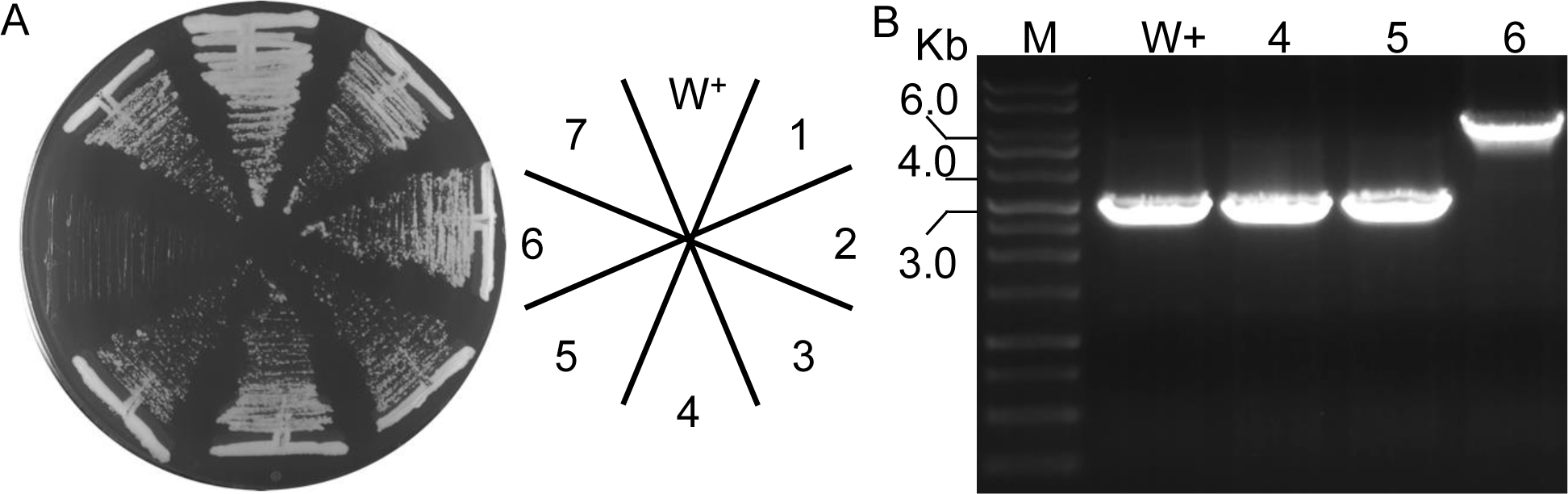
*MDR1* gene deletion in *C. auris*. (A) Wild-type *C.auris* was transformed with a cassette designed to replace *MDR1.* (A) Nourseothricin resistant transformants were transferred to YPD + cyclohexi(i)mide (50μg/ml). (B) Diagnostic PCR using genomic DNA extracted from W+ and three transformants tested in (A) using oligonucleotide primers MdrF2/MdrR3. Sizes of markers (M) are indicated on the left.

A second drug resistance marker gene could not be used for generating a *cdr1Δmdr1Δ* double delete, because *C.auris* is multidrug resistant. Accordingly, the *NAT1* resistance gene was re-cycled by depriving the *cdr1Δ::NAT1-Clox* of methionine and cysteine to induce the Cre-*lox*P removal of *NAT1*. A *cdr1Δ* was isolated as a slow-growing colony from YPD agar containing a sub-lethal concentration of nourseothricin (25 μg/mL). Recombination across the two *lox*P sites removes the *NAT1* cassette and leaves one *lox*P site intact, therefore, the strain is referred to as *cdr1Δ::lox*P from now on. As expected, the wild-type and *cdr1Δ::lox*P strains could not grow in the presence of 200 μg/mL nourseothricin (Figure 3B, sectors 1 and 3), in contrast to the *cdr1Δ::NAT1-Clox* (Figure 3B, sector 2). Additionally, a diagnostic PCR reaction using primers that anneal 5’ and 3’ to the *CDR1* locus confirmed the removal of the *NAT1* cassette in *cdr1Δ::lox*P, giving rise to an expected amplicon of 1905bp in contrast to the 6164bp amplicon from the wild-type (Figure 3C).

**Figure 3.**
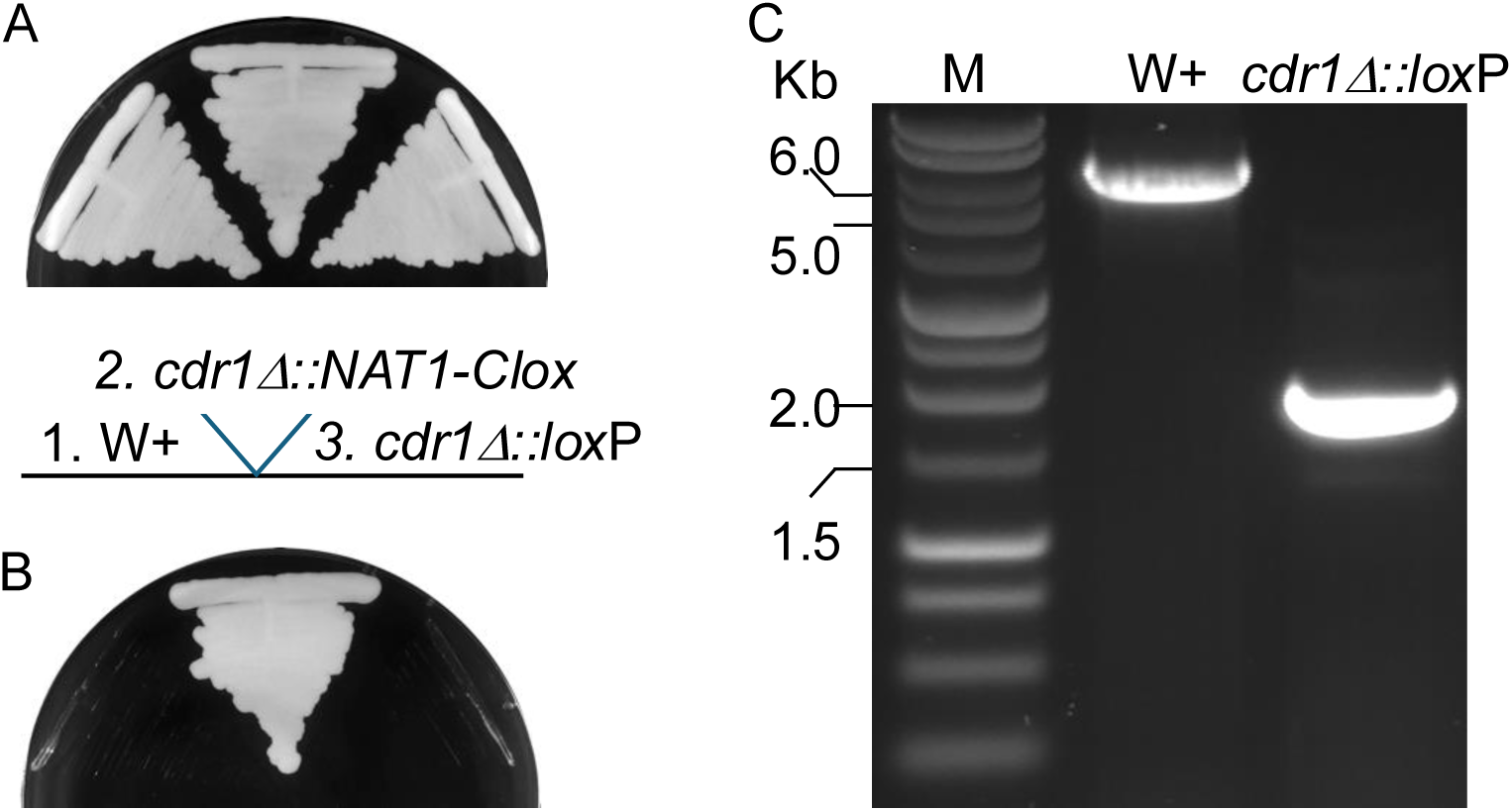
Removal of the *NAT1* drug resistance marker gene from *cdr1Δ::NAT1-Clox.* (A) Cells were grown on YPD with (B) and without (A) 200 μg/mL nourseothricin at 30°C for 5 days. Wild-type (W+) *C. auris* and *cdr1Δ::NAT1-Clox* display nourseothricin sensitivity and resistance, respectively, as expected. *cdr1Δ::lox*P was generated by resolution of the *NAT1* marker gene *via* Cre-mediated recombination across *lox*P sites. (B) Diagnostic PCR using genomic DNA extracted from W+ and *cdr1Δ::lox*P using oligonucleotide primers CDOUTF2/CDOUTR.

*MDR1* was deleted in *cdr1Δ:lox*P by replacing *MDR1* with a targeted deletion cassette. Cells lacking both *CDR1* and *MDR1* are expected to be hypersensitive to cyclohexi(i)mide in comparison to deletion of either drug pump gene alone, and were identified by testing one hundred and twenty nourseothricin resistant transformants on YPD solid agar supplemented with a low concentration of cyclohexi(i)mide (1 μg/mL). Out of the one hundred and twenty transformants, three did not grow after one week of incubation at 30°C (Figure 4 A and B, sectors 1, 2 and 3). An amplicon of 5965bp could be generated from these three strains (Figure 4C, lanes 1, 2 and 3) indicating successful deletion of *MDR1*, in contrast to the 2966bp amplicon corresponding to intact *MDR1*, generated as expected from the W+ and *cdr1Δ::lox*P strains. One of the three cyclohexi(i)mide hypersensitive *cdr1Δ::lox*P *mdr1Δ::NAT1-Clox* strains was selected for further work (and is referred to as *cdr1Δ mdr1Δ* from now on)

**Figure 4.**
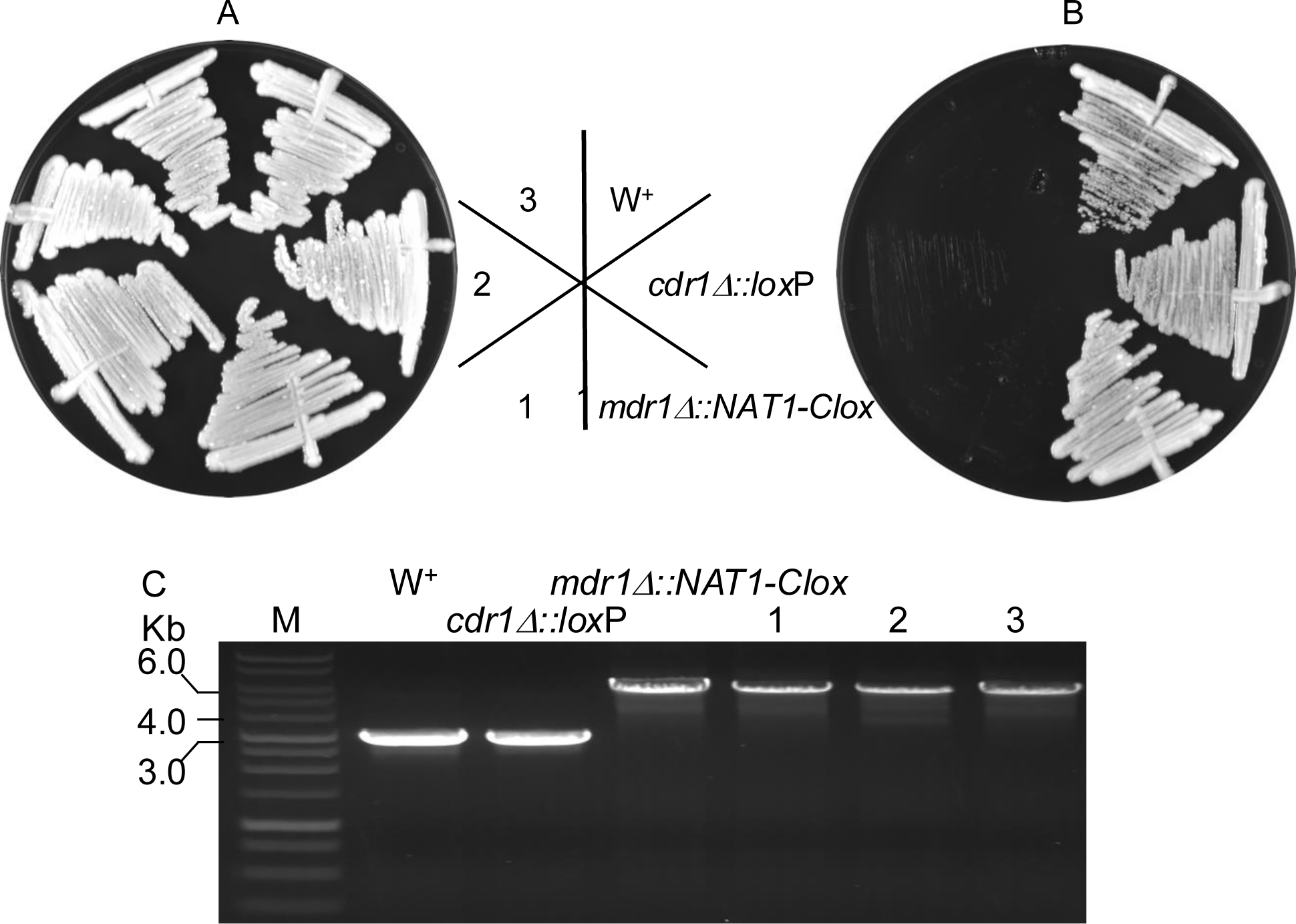
Generating a *cdr1Δmdr1Δ* double delete. The *cdr1Δ:lox*P was transformed with a cassette designed to replace *MDR1*. (B) Of one hundred and twenty nourseothricin resistant colonies, three were hypersensitive to 1 μg/mL cyclohexi(i)mide. Growth of these three strains compared to W+, *cdr1Δ::lox*P, *mdr1Δ::NAT1-Clox* on (A) YPD and (B) YPD+1 μg/mL cyclohexi(i)mide (C) Diagnostic PCR using genomic DNA extracted from the strains indicated in (A) using oligonucleotide primers MdrF2/MdrR3.

### 3.2 Minimum inhibitory concentrations

With increased drug sensitivity already observed for cyclohexi(i)mide with the mutants, particularly for the double mutant strain *cdr1Δmdr1Δ*, the MICs of other commonly used antifungals were also tested(Table 1).

**Table 1.**
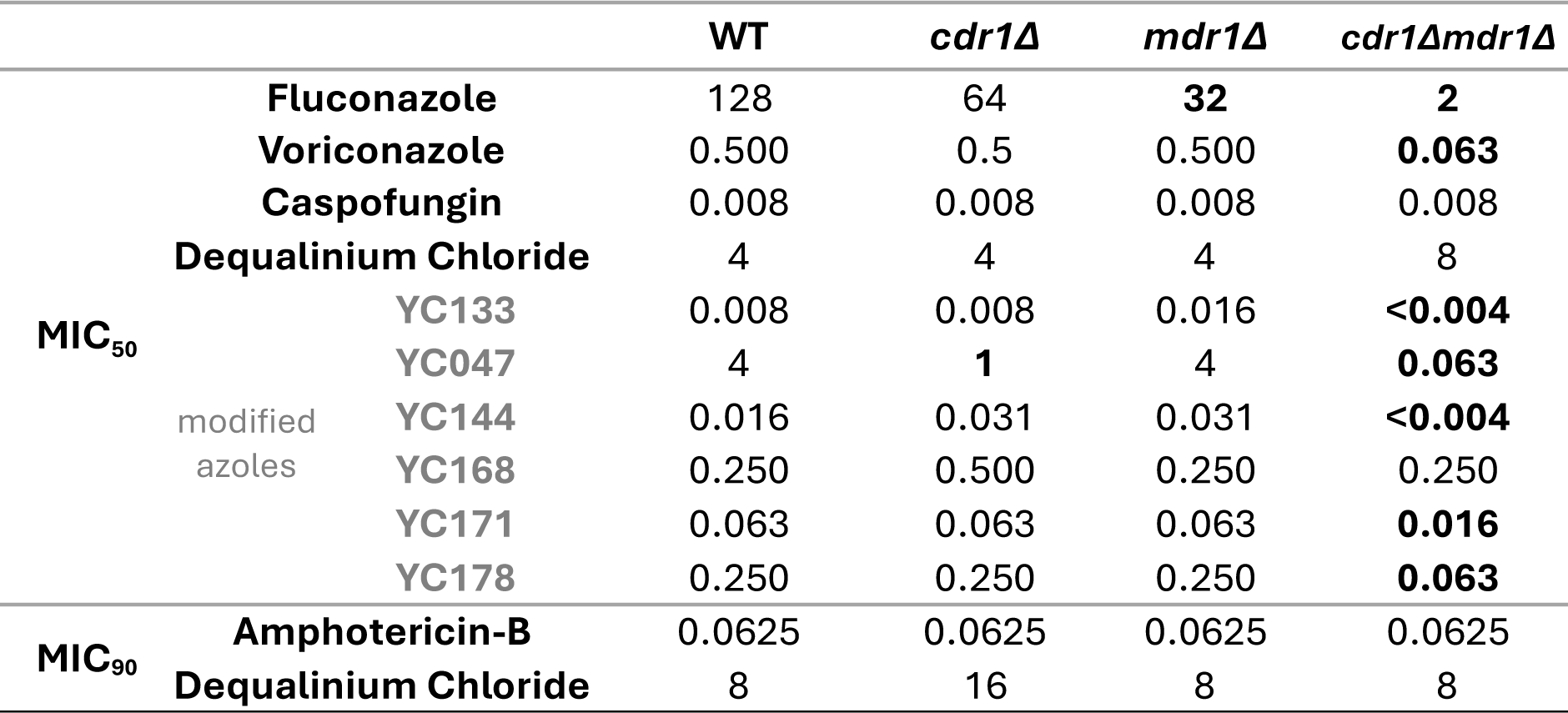
Drug sensitivity in wild-type and mutant strains of *C. auris*. Minimum inhibitory concentrations required to inhibit 50% growth (MIC_50_) and 90% growth (MIC_90_) for clinically used antifungals and a panel of modified azoles, which have been rationally designed to reduce their own efflux. Values in bold indicate a 4-fold or greater change in MIC_50_ relative to wild-type (WT). Measurements are in mg/L.

A 64-fold increase in susceptibility to fluconazole and 8-fold increase in susceptibility to voriconazole was observed for *cdr1Δmdr1Δ*, compared to wild-type *C. auris* (Table 1). Individually deleting each gene had little to moderate increase in susceptibility to fluconazole and voriconazole. This could be a result of compensatory activities between Cdr1 and Mdr1, where one overcomes the absence of the other to continue effluxing of drugs from the cells. In contrast, little to no difference was observed in MIC values between wild-type and any of the mutant strains for the antifungal drugs amphotericin-B (a polyene) caspofungin (an echinocandin) and the antiseptic dequalinium chloride (a quaternary ammonium cation).

A small library of modified azoles, rationally designed to reduce their own efflux through Cdr1 and Mdr1, were also tested against these mutant strains (Table 1). The azole core was modified by the addition of a small pharmacophore, identified through molecular modelling, which interacts with the inhibitor-associated hydrophobic pockets within the efflux pump. This modification introduces efflux pump inhibitory properties into the modified molecule, making them less susceptible to efflux while maintaining interaction with the intracellular target (Laws *et al.*, PCT/GB2018/051468, Rahman *et al*., PCT/GB2021/050268).

The activity of the modified azole YC168 was the same across the wild-type and mutant strains. This compound bypasses the Cdr1/Mdr1 mediated efflux pathway, indicating that the fragment that modified the azole has an effect that is confined to the drug pumps. In contrast, the remaining modified azoles were clearly still being impacted by a Cdr1/Mdr1 mediated efflux pathway, albeit to different extents, ranging from a ca. 63- fold increase in susceptibility of *cdr1Δmdr1Δ* compared to wild-type for YC047, to a ca. 4-fold increase for YC178. This suggests that the action of the fragments that were used for the modification of azoles was not confined to the inhibition of efflux pumps and instead has an additional role, i.e. an off-target effect, demonstrating the utility of the double knockout mutant strain in screening and validating the development new antifungal agents that can overcome efflux.

### 3.3 *Galleria mellonella* infections (virulence)

The *Galleria mellonella* infection model was utilised to assess whether knockout of the efflux pumps impacted virulence of the strains (Figure 5). The survival curves showed no statistically significant difference between wild-type *C. auris*, the single *cdr1Δ* knockout or the double knockout *cdr1Δmdr1Δ*. Although a statistically significant difference was observed between *mdr1Δ* and wild-type, the difference itself was very small (p≤0.05).

**Figure 5.**
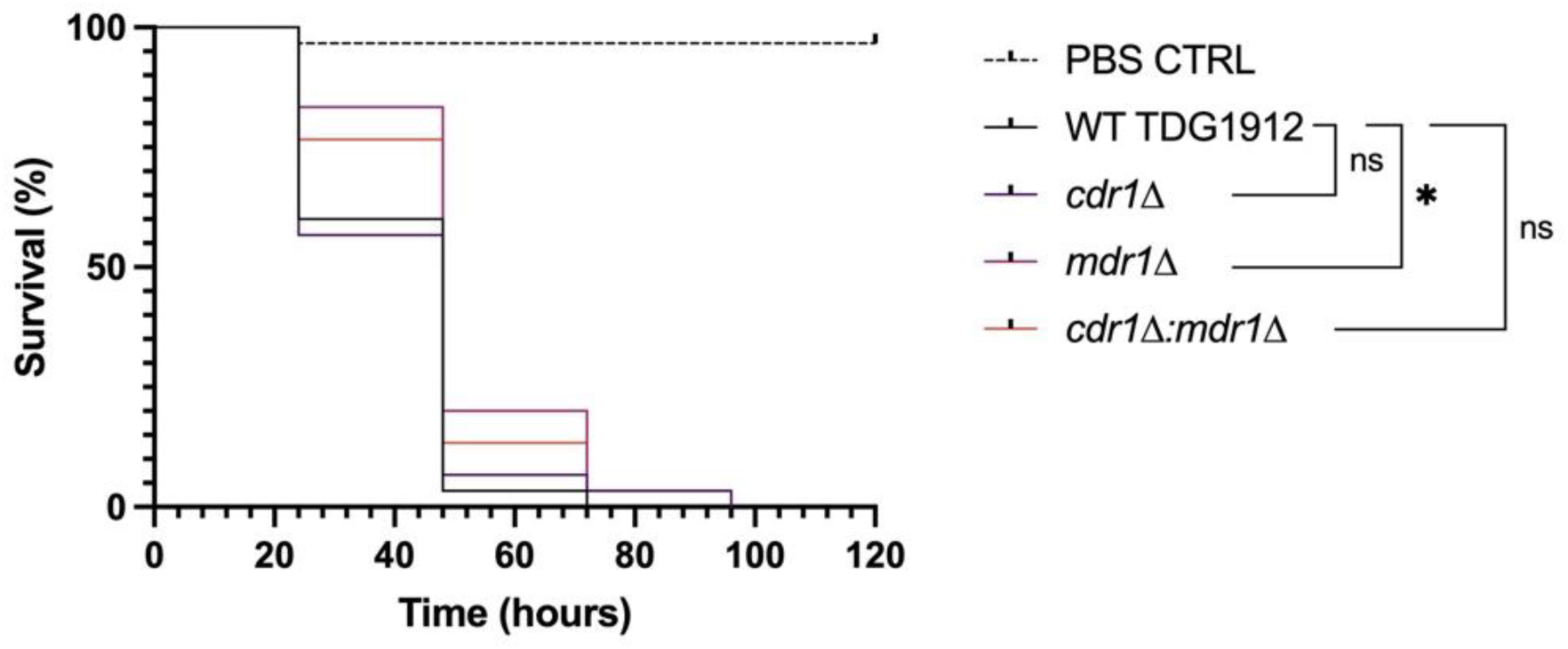
Comparing wild-type *C. auris* and *cdr1Δ*, *mdr1Δ* and *cdr1Δmdr1Δ* virulence. *G. mellonella* infection survival curves were plotted to assess virulence of the wild-type and mutant strains. A statistically significant difference was observed between wild-type and *mdr1Δ.* No statistically significant difference was observed between wild-type and *cdr1Δ,* and *cdr1Δmdr1Δ,* as calculated by log rank (Mantel-Cox) test (n=30, * = p≤0.05, ns = p>0.05).

### 3.4 Biofilm formation

The impact of knocking out the efflux pumps on biofilm formation was analysed using two assays; crystal violet staining (biofilm biomass) and XTT-colorimetric assay (biofilm viability). Crystal violet staining of biofilm formation (represented as % staining of the wild-type) after 24 hours showed a modest but statistically significant difference between wild-type and each of the transporter pump mutants, *cdr1Δ*, *mdr1Δ* and *cdr1Δmdr1Δ*, with the greatest difference observed between *cdr1Δmdr1Δ* and wild-type (Figure 6A). Compared to wild-type, biofilm biomass reduced to 86.6% (*SD*: 5.799, *SEM*: ± 1.450), 90.8% (*SD*: 5.695, *SEM*: ± 1.424) and 69.9% (*SD*: 5.748, *SEM*: ± 1.437) for *cdr1Δ*, *mdr1Δ* and *cdr1Δmdr1Δ*, respectively. In comparison, little to no statistically significant difference was observed in XTT-metabolism assessed biofilm formation (represented as % staining of the wild-type) after 48 hours (Figure 6B).Biofilm formation values for *cdr1Δ*, m*dr1Δ* and *cdr1Δ mdr1Δ*, compared to wild-type, were 111.6% (*SD*: 13.52, *SEM*: ± 3.379), 97.9% (*SD*: 12.66, *SEM*: ± 3.166) and 99.0% (*SD*: 13.62, *SEM*: ± 3.404), respectively.

**Figure 6.**
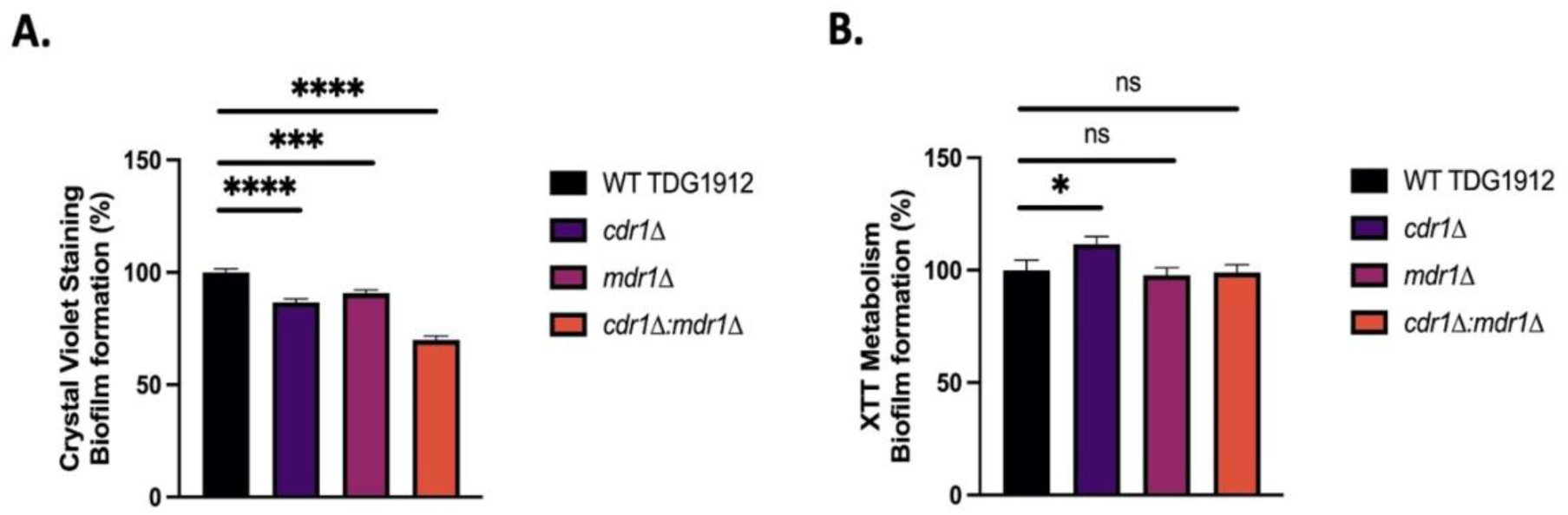
Comparing wild-type *C. auris* and *cdr1Δ, mdr1Δ* and *cdr1Δmdr1Δ* biofilm formation. (A) Crystal violet staining was measured (at OD_570_) of biofilms formed by each strain after 24 hours in YPD broth, at 37°C. A statistically significant difference was observed between wild-type (WT) and all three mutant strains (*cdr1Δ, mdr1Δ,* and *cdr1Δmdr1Δ*) as calculated by T-test (n=48, *** = p≤0.001, **** = p≤0.0001). (B) XTT metabolism, after 3 hours, was measured of biofilms formed by each strain after 48 hours in RPMI. A statistically significant difference was observed between wild-type and *cdr1Δ*. No statistically significant difference was observed between wild-type (WT) and *mdr1Δ*, *and cdr1Δmdr1Δ*, as calculated by T-test (n=48, * = p≤0.05, ns = p>0.05).

We generated the *cdr1Δmdr1Δ* double knockout in order to explore the possibility of using a strain with compromised levels of efflux as a tool to facilitate drug discovery. Rapidly growing yeast cells are generally more susceptible to drugs, due to their active metabolic and biosynthetic processes. Accordingly yeast strains that grow slowly are less affected by drugs because these pathways are less active. Complications arising from negative or positive correlations between growth rate and drug sensitivity are not an issue when using the *cdr1Δmdr1Δ* double knockout, because its growth dynamics are similar to those displayed by the wild-type (Figure 7).

**Figure 7.**
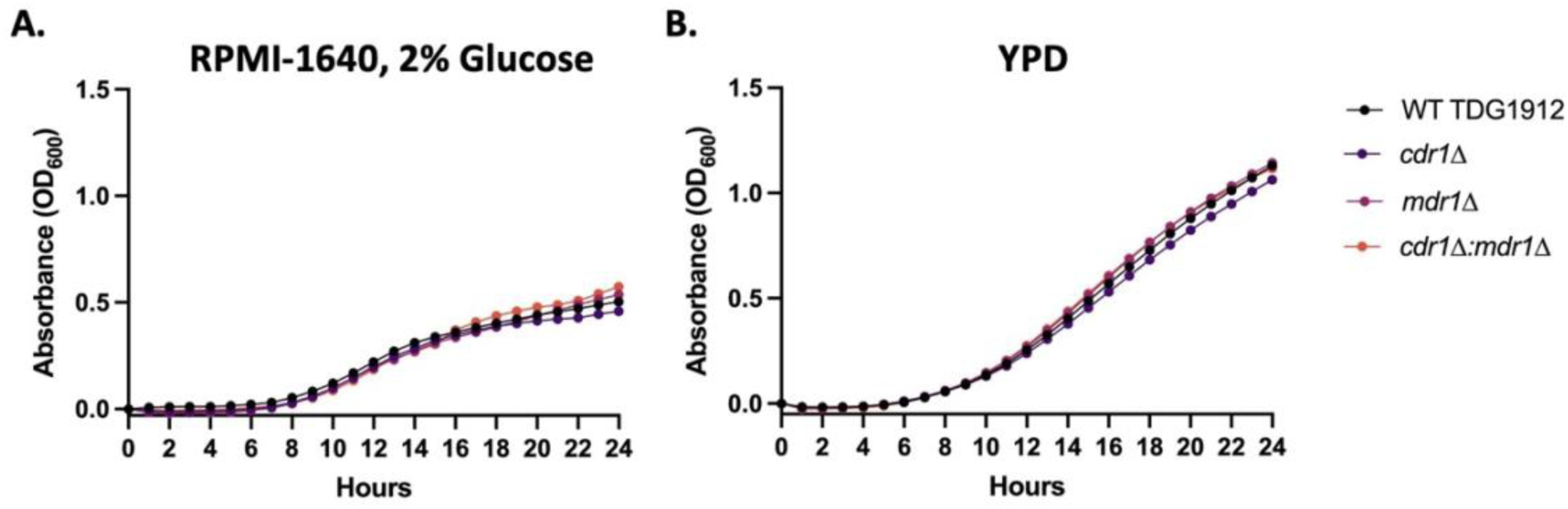
Growth dynamics for wild-type *C. auris* and the mutant strains *cdr1Δ, mdr1Δ* and *cdr1Δmdr1Δ.* (A) Growth curve for strains in RPMI-1640 broth, supplemented with 2% glucose. (B) Growth curve for strains in YPD broth. (n = 9 for each strain in each experimental condition).

### 3.5 High-throughput screening of drug-like compounds

To demonstrate the utility of the double knockout strain for screening compound libraries for candidate scaffolds for drug discovery, a library of 2000 small drug-like compounds was tested at a fixed concentration of 10 µM against the wild-type and *cdr1Δmdr1Δ C. auris* strains (Figure 8). A ‘hit’ was defined as any compound which inhibited growth of either strain by 50% or more. Out of the 2000 drug-like compounds, eleven compounds were identified which inhibited growth of the double knockout *cdr1Δmdr1Δ* strain by 50% or more, three of which resulted in >90% growth inhibition. Only four of these compounds also inhibited growth of the wild-type strain by 50% or more, one of which resulted in >90% growth inhibition. This clearly demonstrates that removing one of the common resistance mechanisms in *C. auris* from the strain used for drug screening can identify a higher number of hit compounds for medicinal drug development projects.

**Figure 8.**
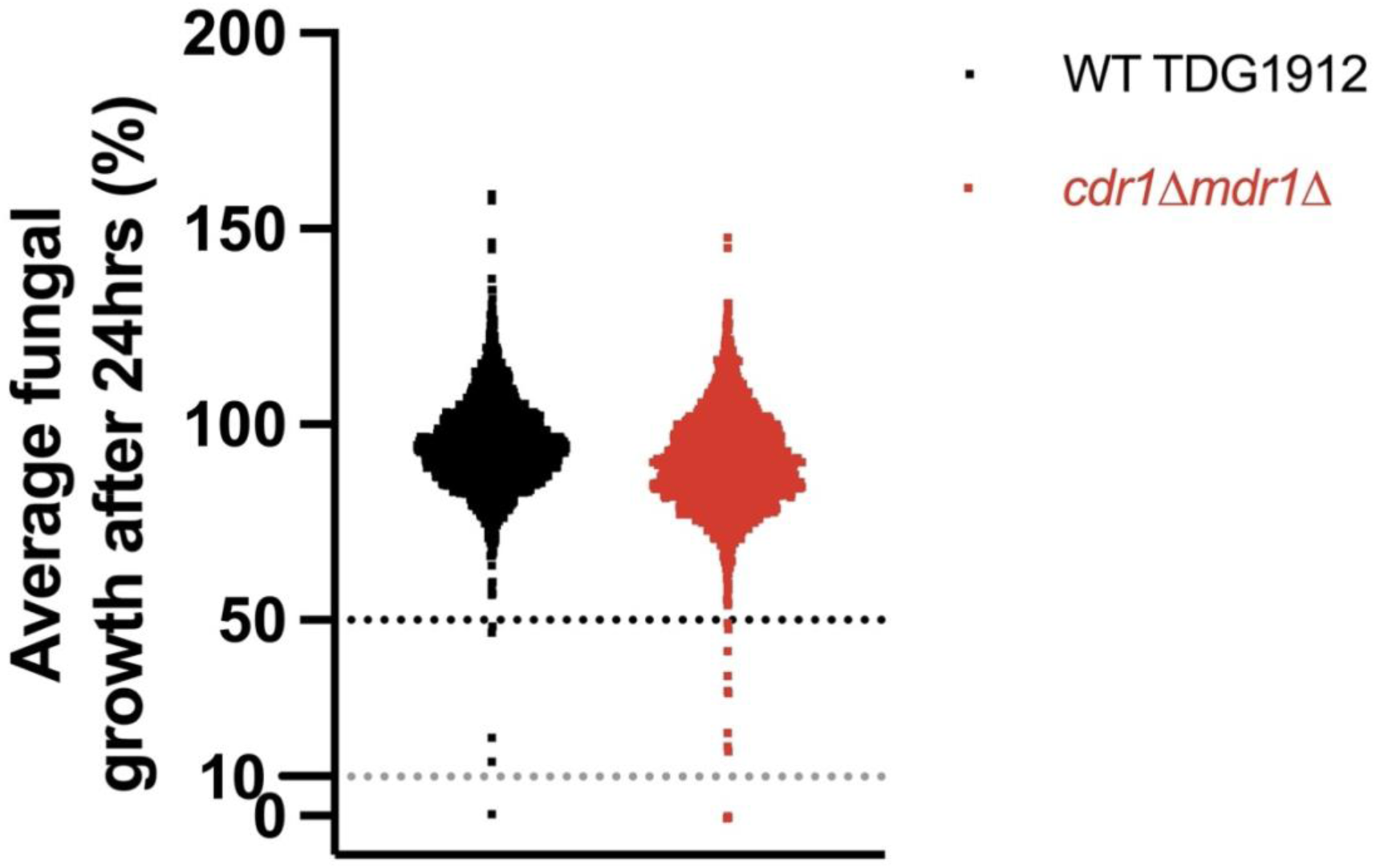
Screening against a 2,000 compound library (Primscreen2, OTAVAChemicals https://otavachemicals.com/sdf). Plot of Average percent fungal growth of *C. auris* wild-type and *cdr1Δmdr1Δ* after 24 hours treatment with 10 μM of each compound. Single dot = Average data for 1 compound. (n=2 per compound).

## 4. Discussion

We have created and characterised, for the first time in the literature, a double efflux pump knockout strain of *C. auris* and used it to identify an increased number of drug candidates compared to screening against a wild-type *C. auris* strain.

Parameters of clinical interest affected by two efflux pumps could be assessed in single and double knockouts. Specifically, little to no difference was observed in virulence in a *G. mellonella* infection model, suggesting that *CDR1* and *MDR1* deletion may not be directly impacting the virulence capacity of *C. auris* cells. Similarly, there was little to no statistically significant difference observed in XTT-metabolism, and therefore biofilm formation, in wild-type *C. auris* and transporter pump mutants. However, results of the crystal violet staining assay indicate a statistically significant difference in biofilm mass, between wild-type, and *cdr1Δ*, *mdr1Δ* and *cdr1Δmdr1Δ* mutants, with the greatest difference observed against the double delete strain. A possible reason for the differences in the results between the two assays could be how each method quantifies biofilm formation. Since crystal violet stains by electrostatically interacting with components of the cell membrane, and transporter pumps are components of the cell membrane, this could impact the overall quantification. Deleting *CDR1* and *MDR1* genes would reduce the amount of protein in the plasma membrane and there may be other secondary effects on the cell surface. This could explain why a small reduction was seen in biofilm formation (%) in strains where either *CDR1* or *MDR1* was deleted, but a slightly larger reduction was seen in the mutant with both genes deleted. Since the XTT-metabolism assay relies on cellular activity, this difference was not seen as the assay was quantifying the biofilm’s capacity for reducing XTT, which evidently was not impacted by the lack of *CDR1* and *MDR1*.

Azoles are known substrates of both *CDR1* and *MDR1,* and testing against azoles highlighted the increased azole sensitivity exhibited by the double delete strain *cdr1Δmdr1Δ*, compared to wild-type *C. auris*. The drug sensitised mutant displayed a 64-fold and 8-fold increase in sensitivity to fluconazole and voriconazole, respectively. This was not observed in the single mutants, demonstrating the redundancy between the Cdr1 and Mdr1 pumps, and the importance of knocking out both of the corresponding genes, for drug discovery. Not surprisingly, a significant difference was not observed between wild-type and *cdr1Δmdr1Δ* against drugs which are not known to be substrates for *Cdr1* or *Mdr1*, namely amphotericin-B, caspofungin and dequalinium chloride.

The advantage of using this strain in drug discovery was confirmed through screening of a large-scale drug-like compound library where an additional seven compounds with antifungal activity were identified against *cdr1Δmdr1Δ*, on top of the four already identified against wild-type *C. auris*. Development of more potent derivatives as well as analysis of the mechanism of action of the compounds, could result in identification of novel antifungal drugs. Using the *cdr1Δmdr1Δ* double knockout can improve the discoverability of novel antifungals, and could be exploited as a drug-discovery tool.

## Ethical approval

Not required.

## Author contributions

Article design and writing: M.H., C.H., J.M.S. and B.P.

Strain construction: M.H.

Drug testing, biofilm and virulence assay: M.H and J. F-P.

Synthesis of modified azole compounds Y.C. and K.M.R.

All authors re-viewed and approved the final version of the manuscript.

## Acknowledgements

Mahnoor Hassan was the recipient of a BBSRC/UKHSA LIDo iCASE doctoral training grant (studentship number BB/M009513/1). UKHSA would like to thank Grant in Aid funding for the Open Innovation Platform (project 113361 and 111742). We thank Al Brown (Exeter University, United Kingdom) for the kind gift of the *NAT1*-Clox plasmid.

## Supplementary file S1

### Constriction of vectors bearing cassettes for gene deletion

#### CDR1

**1. Amplification of domain 5’ to *CDR1***

**Template: *C.auris* genomic DNA**

**HIFI_CdrF1**

<u>CGGCCCACGTGGCCACTAGTACTTC</u>ACCCAGCGAATCTGTAAGCTAAAAG

Overlap with pSL1183 underlined

**CloCdrRev**

<u>TATTACAATTCACTGGCCGT</u>GGAGATGGAAAAGTGAAGATTTGCCTT

Overlap with NAT-Clox PCR fragment underlined

**2. Amplification of *Nat1-Clox*** (composed of *Nat1*, plus *Cre* recombinase gene under the control of the inducible *MET3* promoter, with a *loxP* site at both ends of the cassette).

Template: pLNMLC [1].

**CLoxF**

ACGGCCAGTGAATTGTAATA

**CloxR**

TCGGAATTAACCCTCACTAA

**3. Amplification of domain 3’ to *CDR1***

Template: *C.auris* genomic DNA **CloCdrFor**

<u>TTAGTGAGGGTTAATTCCGA</u>ATTATTGATTATTAGATTGAATACCAT

Overlap with NAT-Clox PCR fragment underlined

**HIFI_CdrR2**

<u>CCTAATTGCAATGATCATCATGACA</u>AGATACTCATCTGAATCACCGTATG

Overlap with pSL1183 underlined

Amplicons from 1,2 and 3 above were assembled into *Xho*1/*Bgl*II digested pSL1183 to give vector pCLOX-CDR1

pSL1183 is a modified version of vector pSL1180 that bears a polylinker composed of restriction enzyme recognition sequences [2]. pSL1183 contains an additional *Asc*1 site between the *BglI*I and *Bam*H1 sites.

**4. Amplification of *CDR1* deletion cassette**

Template: pCLOX-CDR1

**CdrF1**

AACTTGTCGGGCAACTCTCATCTTC

**CdrR2**

AGATACTCATCTGAATCACCGTATG

#### MDR1

**5. Amplification of domain 5’ to *MDR1***

Template: *C.auris* genomic DNA

**HIFI_MdrF1**

<u>CGGCCCACGTGGCCACTAGTACTTC</u>TTCGTACGGTCCTGTCAATATGCAG

Overlap with pSL1183 underlined

**CloMdrRev**

<u>TATTACAATTCACTGGCCGT</u>GTGGAGATTGAAGATGCGTTGAAATTT

Overlap with NAT-Clox PCR fragment underlined

**6. Amplification of domain 3’ to *MDR1***

Template: *C.auris* genomic DNA

**CloMdrFor**

<u>TTAGTGAGGGTTAATTCCGA</u>TAGATTGATTGATACATTTCTTTTACA

Overlap with NAT-Clox PCR fragment underlined

**HIFI_MdrR2**

<u>CCTAATTGCAATGATCATCATGACA</u>CTTCACTGATGCTTGTACATATGGC

Overlap with pSL1183 underlined

Amplicons from 5,2 and 6 above were assembled into *Xho*1/*Bgl*II digested pSL1183 to give vector pCLOX-MDR1

7. Amplification of *MDR1* deletion cassette

Template: pCLOX-MDR1

**MdrF1**

TTCGTACGGTCCTGTCAATATGCAG

**MdrR2**

CTTCACTGATGCTTGTACATATGGC

